# Overcome Competitive Exclusion in Ecosystems

**DOI:** 10.1101/312645

**Authors:** Xin Wang, Yang-Yu Liu

## Abstract

Explaining biodiversity in nature is a fundamental problem in ecology. One great challenge is embodied in the so-called competitive exclusion principle^1-4^: the number of species in steady coexistence cannot exceed the number of resources^4-7^. In the past five decades, various mechanisms have been proposed to overcome the limit on diversity set by the competitive exclusion principle^8-25^. Yet, none of the existing mechanisms can generically overcome competitive exclusion at steady state^4,26^. Here we show that by forming chasing triplets in the predation process among the consumers and resources, the number of coexisting species of consumers can exceed that of resources at steady state, naturally breaking the competitive exclusion principle. Our model can be broadly applicable to explain the biodiversity of many consumer-resource ecosystems and deepen our understanding of biodiversity in nature.

---

In Darwin’s theory of evolution^27^, survival of the fittest, i.e., the less competitive species dies out, implicates the notion of competition exclusion. In 1928, Volterra illustrated^3^ mathematically that when two species competing for a single resource, one competitor must die out except that if the hunting to death rate ratio is exactly the same for the two competing species (a Lebesgue zero-measure set for parameters). Those results were absorbed in the competition exclusion principle (CEP) formulated in 1960s^4-7^. The CEP can be mathematically described in the consumer-resource model framework. Consider *M* types of consumer species competing for *N* types of resources. Each consumer can feed on one or multiple types of resources. Consumers do not directly interact with each other via other mechanisms except the competition for the resources. According to the CEP^4-7^, at steady state the number of coexisting species of consumers cannot exceed that of resources, i.e. *M ≤ N*.

The classical proof^5-7^ of the CEP is demonstrated in Fig. 1. Consider the simplest case *M =* 2, *N =* 1, i.e. two species *C*_1_ and *C*_2_ competing for a single resource *R*. The generic population dynamics of this consumer-resource ecosystem is shown in Fig. 1b. At steady state, if the two species coexist, we have *f_i_* (*R*)/*D_i_ =* 1 (*i* =1, 2). This requires that the two curves *y = f_i_ (R*)/*D_i_* (*i* =1, 2) should cross the line *y* =1 at the same point, which is typically impossible (Fig. 1c), unless the model parameters satisfy certain constraint. Similar proof strategy applies to the case of *M =* 3, *N = 2* (see Fig. 1e-h). In fact, we can extend the proof to any positive *N* and *M*^5^.

**Figure 1.**
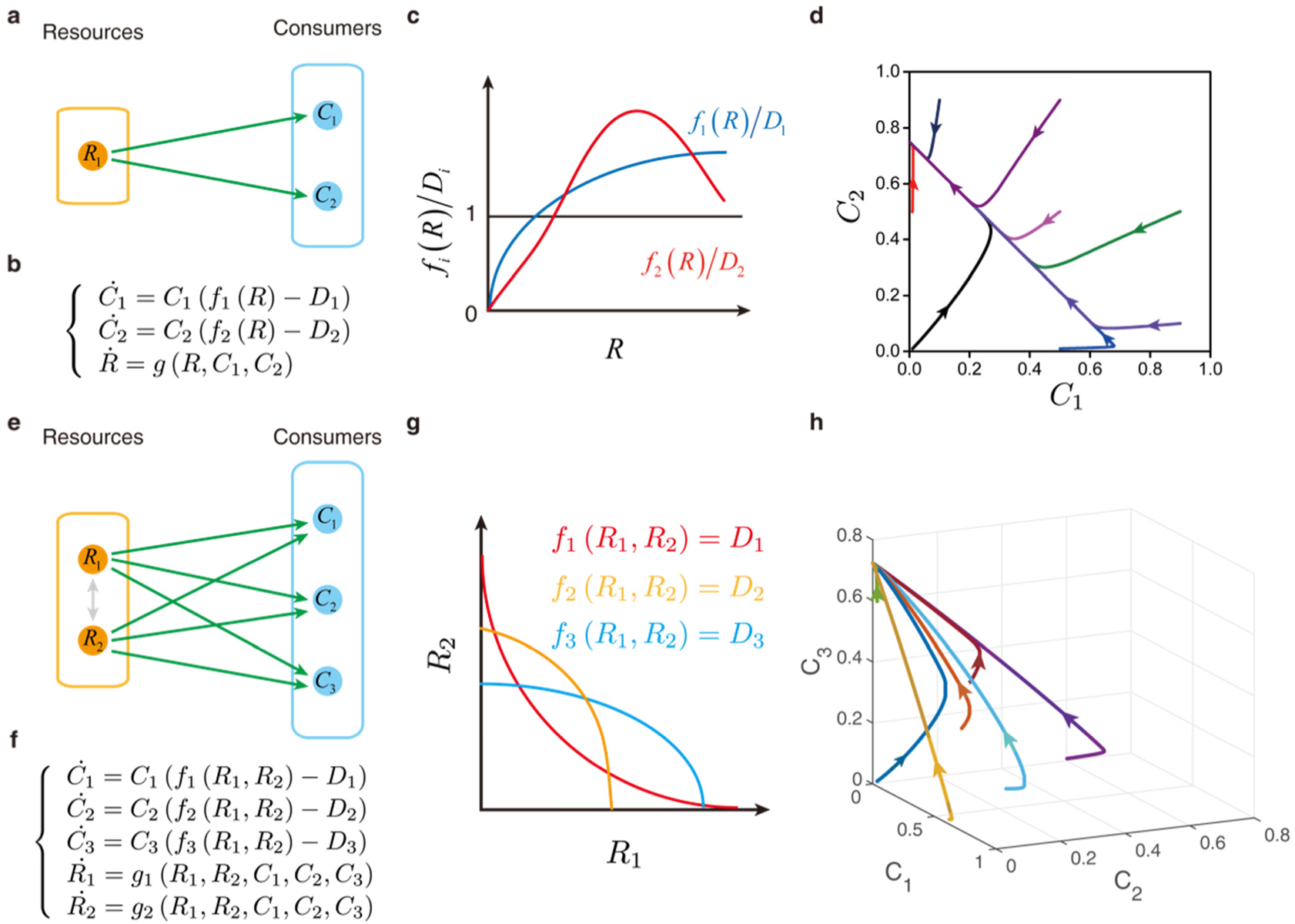
Classical proof of the competitive exclusion principle. (a) Bipartite graph of the scenario of *M =* 2, *N* = 1. (b) Generalize mathematical description of the models in the classical proof, where *f_i_* (*i* =1, 2), *g* can be any function of the required form, *D_i_* (*i* =1, 2) stand for mortality rate of the consumers. (c) At steady state, if all consumer species coexist, *f_i_* (*R*)/*D_i_ =* 1 (*i* =1, 2). This requires that three lines *y = f_i_* (*R*)*/D_i_* (*i* =1, 2) and *y* =1 own a common node, which normally cannot happen. (d) Representative phase portrait of the trajectories, two consumer species cannot coexist when *N* =1. Here *f_i_* (*R*) = 0.002*R* (*i* =1, 2); *g* (*R*, *C*_1_, *C*_2_) = *R* (1 − *R − C*_1_ − *C*_2_); *D*_1_ *=* 0.0006, *D*_2_ = 0.0005. All trajectories start at *R =* 0.01. (e) Bipartite graph of the scenario of *M =* 3, *N* = 2. (f) Generalize mathematical description of the models in the classical proof, where *f_i_*, *g_i_* (*i* =1, 2, 3; *j* =1, 2) can be any functions of the form, *D_i_* (*i* = 1, 2, 3) denote for mortality rate of the consumers. (g) The relations between two species of resources. At steady state, if all consumer species coexist, *f_i_* (*R*_1_*,R*_2_) = *D_i_* (*i* =1, 2, 3). Generically, three curves would not cross at exactly the same point, hence the three consumer species cannot coexist at steady state. (h) Representative phase portrait of the trajectories; three consumer species cannot all coexist at steady state. Here 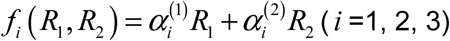, 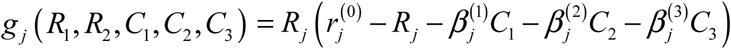 (*j* = 1,2); *D*_1_ = 0.0006, *D*_2_ = 0.0005, *D*_3_ = 0.0004, 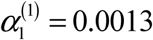, 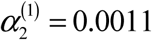, 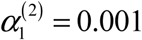, 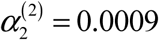, 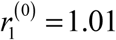, 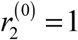, 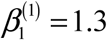, 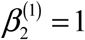, 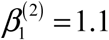, 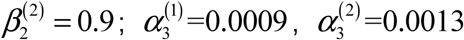, 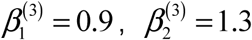. In the initial condition, *R*_1_ = 0.01, and *R*_2_ = 0.01 for all trajectories.

In the classical CEP framework there are only two scenarios that permit *M > N*: (i) consumers’ densities never reach steady state: they may fluctuate consistently^20,21^ or be in a chaos^21,22^; (ii) some pathological cases with zero measure that occur when the system parameters satisfy certain accidental constraints^3,23^. It is still unknown if we can break the constraint of CEP generically at steady state, without assuming any special model parameters.

Here, we consider the predation process between the consumers and resources, and assume both are biotic. We explicitly consider that the population structure of consumers and resources: some are wandering around freely, some are chasing each other. When a consumer meets a resource, they form a chasing pair, denoted as *R*^(^*^P^*^)^ ⋁ *C*^(^*^P^*^)^, where the superscript ‘P’ stands for ‘pair’. The resource can escape with probability *d* or be caught and consumed by the consumer with probability *k*. Such a predation kinetics commonly takes a Michaelis-Menten form: 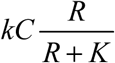, with 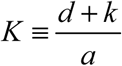, which corresponds to the Holling’s type-II functional response^28^ and is widely adopted in consumer-resource models^20,29^. This form, in fact, agrees with the growth rate function in the classical proof^5-7^, where 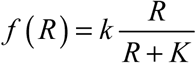. Nevertheless, the Michaelis-Menten kinetics is a good approximation only if the resource population is much larger than the consumer population, i.e., *R* ≫ *C* (see SI Sec.3 for details). When this condition is not satisfied, the growth rate functions follows *f* (*R*, *C*)^30^ rather than *f* (*R*). The C-dependency in the growth rate function makes the classical proof^5-7^ invalid, implying a potential mechanism to break the CEP.

Interestingly, we find that the chasing-pair scenario is still not enough. For example, in case *M =* 2 and *N =* 1 (Fig. 2a), The population dynamics of the system can be described by:

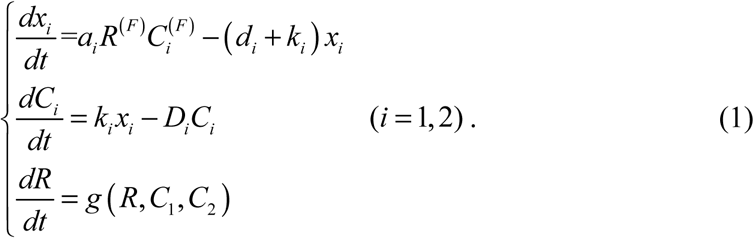

**Figure 2.**
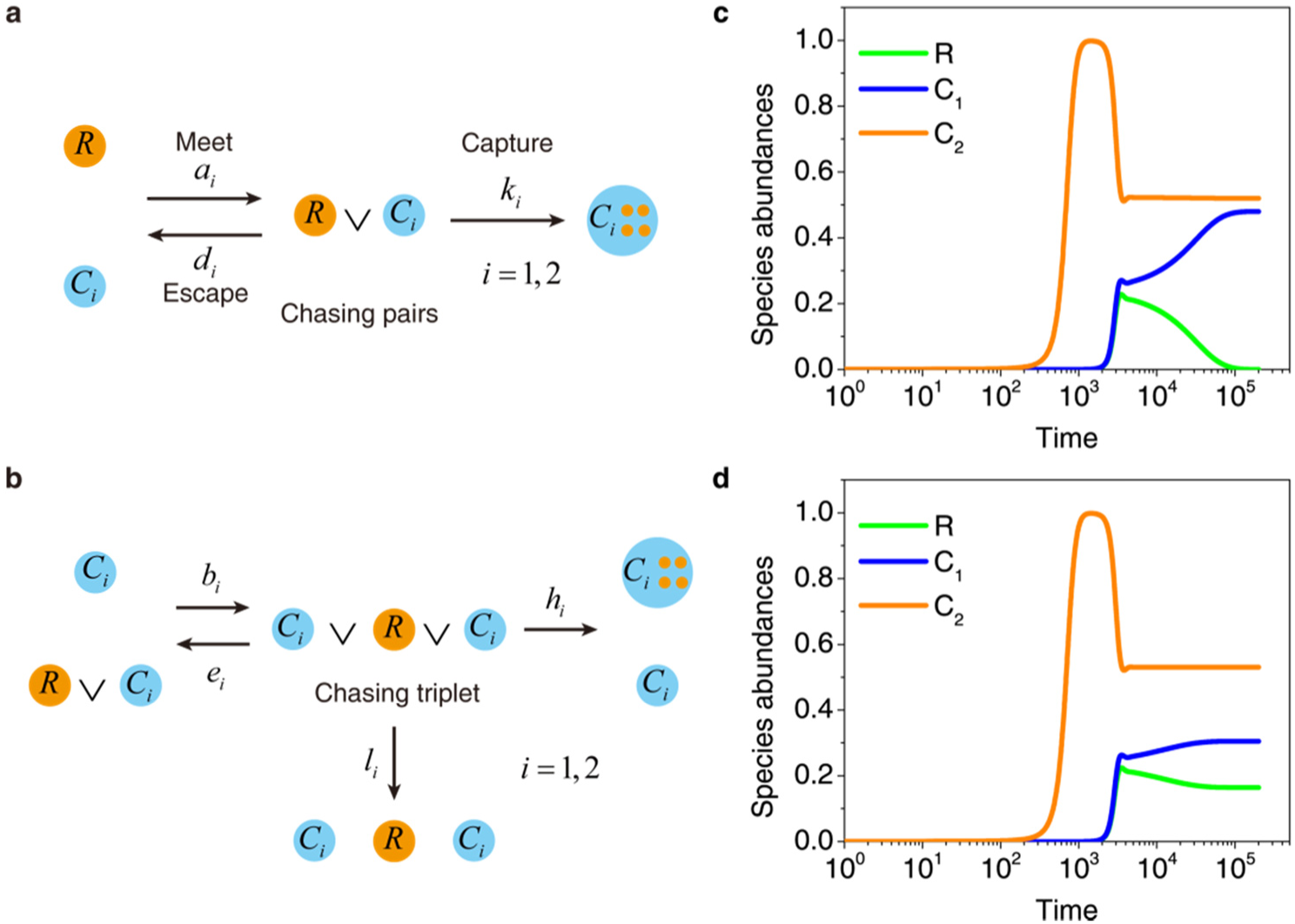
Schematic of the predation process between consumers and resources. *M = 2*, *N =* 1. (a) Formation of a chasing pair between a consumer and a resource. Here *a_i_* is the encounter rate between a consumer and a resource to form a chasing pair, *d_i_* is the escape rate of a resource out of a chasing pair, and *k_i_* is the capture rate of consumers in a chasing pair. (b) Formation of a chasing triplet among two consumers of the same species and a resource. Here *b_i_* is the encounter rate between a consumer and an existing chasing pair to form a chasing triplet, *e_i_* is the escape rate of a consumer out of a chasing triplet, *h_i_* is the capture rate of consumers in a chasing triplet. (c) Time course of the abundances of two consumers (*M* =2) and one resource (*N* =1). In the presence of chasing pairs, consumer species cannot coexist at steady state. (d) Time course of the abundances of *M* = 2 and *N* = 1. In the presence of chasing pairs and chasing triplet, consumer species coexist at steady abundance. Here, we assume that the dynamics of the resources follow the classical MacArthur’s consumer-resource model^31,32^: 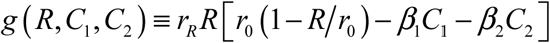. (c) & (d) were simulated from equation 2. In (c): *b_i_ = e_i_ = h_i_ = l_i_ =* 0(*i* =1,2); In (d): *b =* 0.1, *e_i_ = l_i_ =* 0.1, *h_i_* = 0.02 (*i* =1, 2); In (c) & (d): *a_i_ =* 0.1, *d_i_ =* 0.1, *k_i_ =* 0.02 (*i* =1, 2), *D*_1_ *=* 1.01*D*_2_, *D*_2_ *=* 0.005, *r_R_* = 0.01, *r*_0_ = 1 and *β_i_* = 1, the initial abundances of (*R*,*C*_1_,*C*_2_) are (0.001, 0.001, 0.001).

Here consumers and resources that are freely wandering around are denoted as 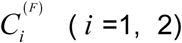 and *R*^(^*^F^*^)^, 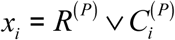 represents the chasing pair, *a_i_* is the encounter rate between a consumer and a resource to form a chasing pair, *d_i_* is the escape rate of a resource out of a chasing pair, and *k_i_* is the capture rate of consumers in a chasing pair. If the two consumers can coexist, we can prove that the steady-state equations yield 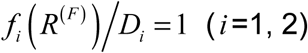, which corresponds to parallel planes in a 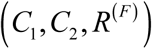 coordinate (Fig. S8b), rendering coexistence impossible (Fig. 2c, see SI Sec.4-5 for details).

To find the mechanism that can generically break the CEP at steady state, we revisit the predation process. We naturally extend the idea of chasing pair to chasing triplet, i.e. two consumers (within the same or from different species) can chase the same resource (Fig. 2b, Fig. S5 & Fig. S6). For example, in case *M =* 2, *N =* 1, a consumer (*C_i_*, *i* =1 or 2) can join an existing chasing pair 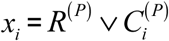 to form a chasing triplet 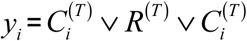 (Fig. 2b). Those consumers and resources that are freely wandering around are denoted as 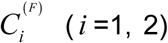 and *R*^(^*^F^*^)^, respectively. The population of consumers *C_i_* (*i* =1, 2) and resources are given by 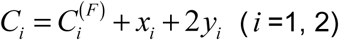 and 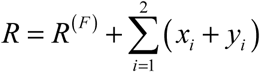, respectively. The population dynamics of the system can be described by (see SI Sec.5 for details):

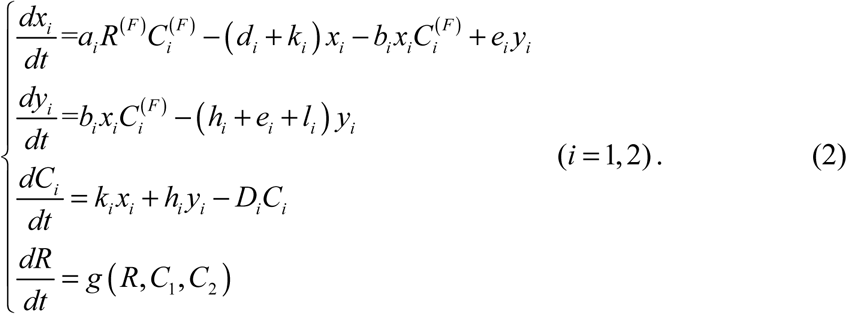

Here *b_i_* is the encounter rate between a consumer and an existing chasing pair to form a chasing triplet, *e_i_* is the escape rate of a consumer out of a chasing triplet, *h_i_* is the capture rate of consumers in a chasing triplet, and *D_i_* (*i* =1, 2) is mortality rate of consumer species *i*.

Note that in the classical model in case *M = 2* and *N = 1* (Fig. 1 a-d), if both consumers species can coexist steadily, the abundance of resources *R* needs to satisfy two equations simultaneously, which is equivalent to require that two parallel planes share a common point, which is impossible (Fig. 4a). In the presence of chasing pairs, as shown in Fig. 4b (See SI Sec.4-5 for details), the requirement for steady coexistence corresponds to parallel surfaces. In the presence of chasing pairs and chasing triplets, as shown in Fig. 4c (See SI Sec.5 for details), the requirement for steady coexistence corresponds to three non-parallel surfaces to cross at one point, which can naturally happen. Hence, the CEP will be broken. Our numerical simulations confirmed this point (see Fig. 2d).

Moreover, we find that the coexisting state can be globally stable (Fig. 3a) as long as the initial abundances of both consumer species are non-zero. And such a globally stable state can exist for a range of parameter values (Fig. 3b). In other words, the broken CEP is not due to a pathological set of model parameters. Note that the violation of CEP in the case of *N =* 1 actually implies that it will be violated for more general cases with *N >* 1 (see SI Sec.6 for details). Note that our model doesn’t allow for unlimited number of consumer species to coexist at steady state for a limited number of resources. Consider the simplest case of *N* = 1, we can show that the parameter space that facilitates steady coexistence shrinks when *M* increases from 1 to 2. We expect that the parameter space would further shrinks with increasing *M*. How does the parameter space shrink is a challenging question that is beyond the scope of the current work.

**Figure 3.**
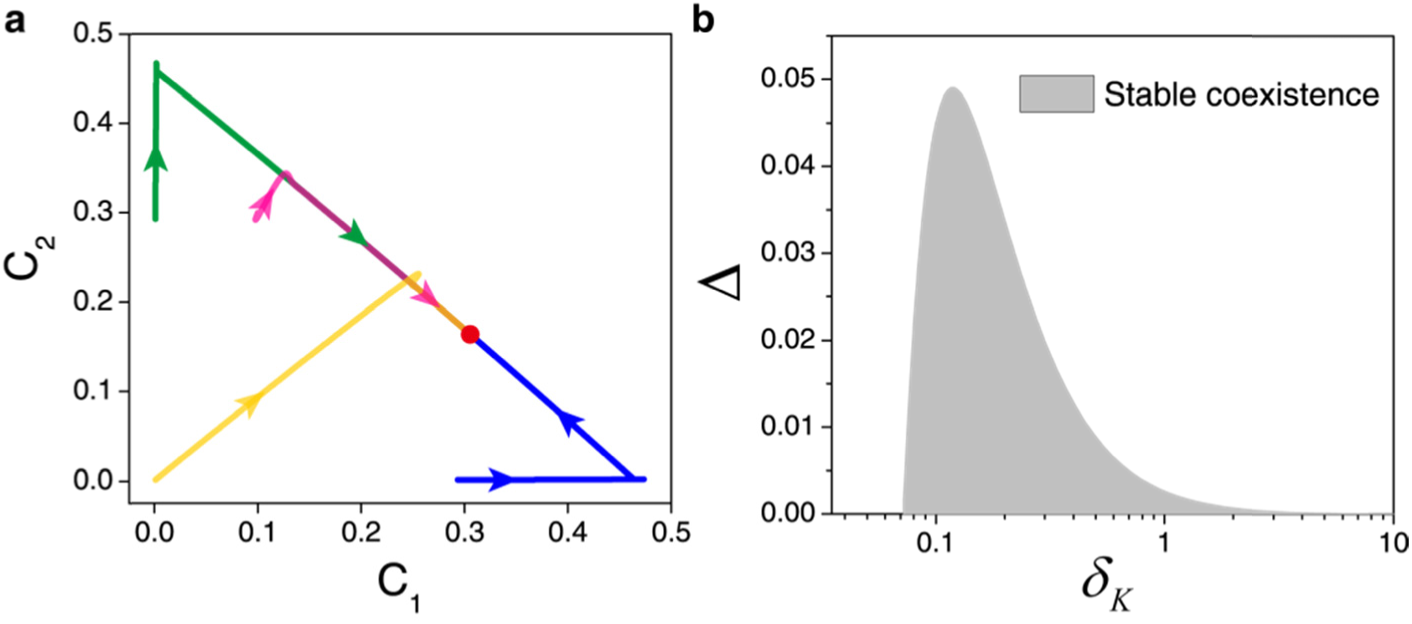
Stable coexistence of two consumer species with one type of resources. (a) Stable attractive property of the co-existing state. Equations and parameters are the same as that in Fig. 2d. (b) The region of stable coexistence (shown in grey) of the parameter set. To measure how large the difference can be tolerated between the two consumer species when achieving stable coexistence, we define Δ = (*D*_1_ − *D*_2_)/*D*_2_, i.e. the relative difference in mortality rate between the two consumer species, and multiply *δ_K_* on *k_i_*, *h_i_*, *d_i_*, *e_i_*, 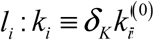, 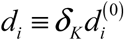, 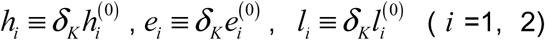. Meanwhile we set *a*_1_ = *a*_2_, *b*_1_ = *b*_2_, 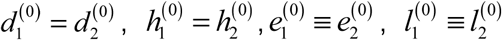, so that the difference between the two species can be reflected on Δ. By varying *δ_K_*, we find that there is upper bound tolerance for difference Δ, below which (the grey region) the two consumer species can stably coexist. The result manifests stable coexistence at non-special model parameters. In (b): *a_i_* = *b_i_* = 0.1, 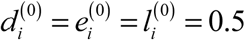, 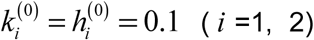, *D*_1_ =(1 + Δ)*D*_2_, *D*_2_ = 0.005, *r_R_* = 0.01, *r*_0_ = 1 and *β_i_* = 1. We have further considered scenarios where chasing triplet is formed between different species of consumers (Fig. S5, combining Fig. S5a & b, See SI Sec.5 for details) or either between or within species (Fig. S6, combining Fig. S6a, b & c, See SI Sec.5 for details). In all cases above, there is a non-special parameter set where the two consumer species can stably coexist (gray region, Fig. S7 b & c).

**Figure 4.**
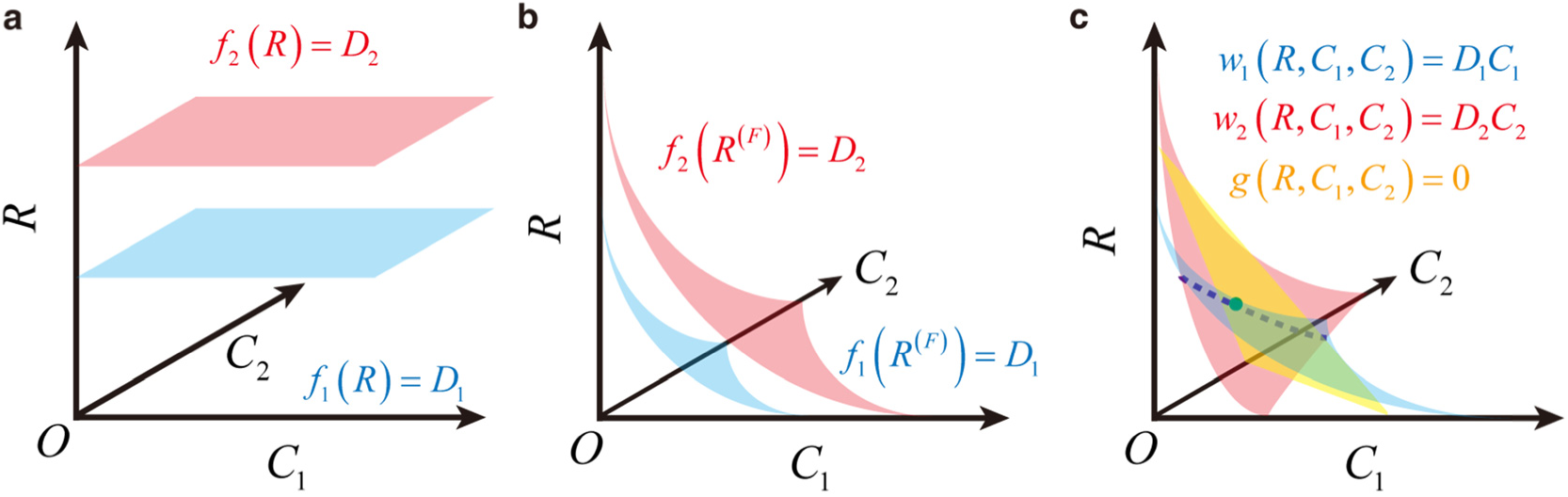
Intuitive explanation of how the formation of chasing triplet liberates the constraint of competitive exclusion. (a) In the classical proof, the red plane and blue plane are parallel to each other, and thus do not have a common point. (b) In the model involving chasing pairs, the red surface and blue surface are parallel to each other, and thus still do not have a common point (see Fig. S8b, SI Sec.4–5 for details) (c) In the model involving chasing pairs and chasing triplets, the three color surfaces (or planes) are not parallel one to another, and thus the red surface and blue surface can own an intersection curve (dashed purple curve), while the three surfaces can intersect at one point (green point) and thus facilitate coexistence. *M =* 2, *N =* 1 in (a) and (b).

The CEP has been proposed for decades. Although it apparently contradicts to the observation of biodiversity in nature, no prior mechanism has challenged this principle at steady state. Here, by taking into account the details of the predation process, especially the possibility to form chasing triplet, we liberate the constraint of competitive exclusion. Given that stable coexistences between species are widely observed or implied in natural ecosystems such as microbiome within human gut, tree species in forests, and planktons in the marine world, the results presented here deepen our understanding on the biodiversity in nature, and may be applicable to broadly diversified consumer-resource systems.

